# Protein Evolution Depends on Multiple Distinct Population Size Parameters

**DOI:** 10.1101/141499

**Authors:** Alexander Platt, Claudia C Weber, David A Liberles

## Abstract

That population size affects the fate of new mutations arising in genomes, modulating both how frequently they arise and how efficiently natural selection is able to filter them, is well established. It is therefore clear that these distinct roles for population size that characterize different processes should affect the evolution of proteins and need to be carefully defined. Empirical evidence is consistent with a role for demography in influencing protein evolution, supporting the idea that functional constraints alone do not determine the composition of coding sequences.

Given that the relationship between population size, mutant fitness and fixation probability has been well characterized, estimating fitness from observed substitutions is well within reach with well-formulated models. Molecular evolution research has, therefore, increasingly begun to leverage concepts from population genetics to quantify the selective effects associated with different classes of mutation. However, in order for this type of analysis to provide meaningful information about the intra- and inter-specific evolution of coding sequences, a clear definition of concepts of population size, what they influence, and how they are best parameterized is essential.

Here, we present an overview of the many distinct concepts that “population size” and “effective population size” may refer to, what they represent for studying proteins, and how this knowledge can be harnessed to produce better specified models of protein evolution.

## Introduction

Understanding how proteins evolve under the influence of natural selection is a central goal of evolutionary biology, as it provides insight into functional constraints and the diversification of genomes. Although it may be tempting to study protein evolution from a predominately biophysical perspective, considering how mutational processes generate variation and population level processes modulate selection is necessary to fully explain extant coding sequences. Counter to the notion that amino acid sequences are determined by functional requirements alone, some of the observed variation is a consequence of the limits of natural selection in finite populations.

The functional synthesis of protein evolution and population genetics has shown that the size of a population (*N*) modulates amino acid sequence divergence as well as the rates and patterns of adaptation [1, 2]. In simple models of population genetics, the standing pool of genetic diversity, the probability of fixation of a new mutation, and the relative fixation probability of a selected mutation all relate to N and have expected values of 2*Nμ*, 1/*N* and 2*Ns*, respectively. Here, s represents the relative selection coefficient of the mutant allele (that is, how fit the mutant is compared to the wild type).

In accord with this view, comparative genomics has linked changes in effective population size to differences in observed properties of proteins, such as stability and other features subject to selection, including binding specificity and avoiding misinteraction [3, 4, 2]. The rate of accumulation of neutral changes is not affected by *N*, as the probability of introduction of a mutation is the inverse of the neutral probability of fixation. However, all changes subject to selective constraint or adaptive pressure are influenced by the population size [5].

Beyond the simplest models, however, the number of individuals in a population is rarely the correct scalar for all of these parameters of interest. It is standard practice, therefore, to employ a variation of it, the *effective* population size (*N_e_*) to account for deviations introduced by any and all of a host of complications such as inbreeding, unequal sex ratios, linked selected sites, population substructure, life-history patterns, or high-variance reproductive strategies [6].

Given its importance in influencing sequence variation, what precisely do we mean when we refer to *N_e_* in the context of protein evolution? There are multiple definitions of effective population size in use. For instance, in population genetics, *N_e_* might be treated as a convenience parameter reflecting the extent of genetic drift (stochastic changes in allele frequency) inferred from a sequence, or as a constant summarizing an unruly history of fluctuating demography or complicated social structure. However, to understand how protein structure and function drive amino acid substitution, we require models that disentangle the factors contributing to neutral and adaptive sequence divergence and describe the underlying biological processes accurately [7, 8]. We refer to these as mechanistic models in this work (though the term is used differently elsewhere [9, 10]).

Here we discuss how to define *N_e_* and how this parameter can be augmented to capture information about mechanistic processes that are distinct from natural selection, a prerequisite to realistically characterizing the evolution of proteins in some scenarios.

## Origins and Historical Applications

The concept of effective population size has its origins in the Wright-Fisher model, which describes the change in allele frequency through time in a single randomly mating population of constant size. This model has a single parameter: the size of the population. *All* predictions from this model could be interpreted as functions of the population size, giving a direct map between any predictable population measurement and a population size that would produce it. Where the model assumptions of Wright-Fisher were applicable, it was reasonable to refer to this parameter in its original sense as the unchanging, haploid, panmictic, neutral, asexual, non-overlapping, population size *N*.

This model proved to be both tractable and powerful for deriving many important properties of evolving gene pools such as quantifying genetic drift, the probabilities of allele loss and fixation, and allele sojourn times within specified frequency ranges. Real populations, and even most interesting population models, however, violate many of the restrictive assumptions made in the Wright-Fisher model. Deviations include population size change, population structure, organisms divided into sexes, selection acting both directly on individual loci and indirectly on linked neutral loci (which we may not wish to include as part of *N_e_*), and non-overlapping generations. Still, for any observation predictable under the Wright-Fisher model, it is possible to ask what population size in the Wright-Fisher model maps to the observed value in the more complicated model. This parameter was given the name ‘‘effective population size”, while the actual size of the population is designated as the census population size or *N_c_*.

Defined this way there is no requirement that the effective population size need be the same for any two different observations from the same population. In a Wright-Fisher population there is only a single value of *N* that is used to make all predictions about the behavior of the model. In a non Wright-Fisher population, the *N_e_* that corresponds to the probability of fixation of a newly arising allele might not be the same value as the *N_e_* that describes, for example, the rate of change in frequency that said allele exhibits or the probability that two randomly sampled individuals share a recent common ancestor. In each case, one must independently compare the observed value to a corresponding Wright-Fisher model, and it is always necessary when talking about the effective size of a population to qualify it with what observation is being described. Careful usage, then, was to explicitly label an effective population size as to what it was describing [11]. Common variants included the inbreeding effective population size (describing the probability that two individuals shared a common ancestor in the previous generation), the variance effective population size (describing the variance in reproductive success among individuals), and the eigenvalue effective population size (describing the leading non-zero eigenvalue of the allele frequency transition matrix). These concepts (like inbreeding structure or variable reproductive success in the population over a lineage of a phylogenetic tree) are directly related to individual protein-specific selective pressures and probabilities of amino acid fixation through the effects of the broader population acting on all proteins.

The more recent introduction of another useful effective population size, the coalescent effective population size, has gone a great way in elucidating the underlying mechanism of the frequent similarities of effective population sizes as well as when more subtle distinctions are required [12, 13, 14, 15]. Many parameters of interest to a population geneticist can be described as properties of the genealogy of a random sample of individuals. Furthermore, many different population models converge to a common underlying form of this genealogy (often referred to as Kingman’s coalescent), each differing only in a single scaling parameter. This includes models with deviations from Wright-Fisher such as unequal sex ratios, structured life-stages, or uniformly elevated or depressed reproductive variance. For population models that converge to Kingman’s coalescent, this scaling factor serves the purpose of the effective population size for *all* of the population genetic statistics determined by sample genealogy. This coalescent effective population size is sometimes defined as the ratio of the expected generation time to the expected time since common ancestry of two homologous chromosomes. It is compelling both as an explanation of why a great many definitions of effective population size often produce very similar numbers, as they are derived from a common rescaling of the underlying genealogy, and as an illustration of when there is *no* single parameter that can meaningfully compare the general behavior of a particular population to one evolving under Wright-Fisher rules. As illustrated in Figure 1, subdivided populations offer a particularly clear example of this type of violation. For any large, random sample of individuals, the genealogy will have an excess of short tips (individuals in small sub-populations sharing recent common ancestry with each other) and longer internal branches (individuals in different sub-populations sharing only distant ancestors) than described by the Kingman coalescent. As the distortions in different parts of the tree are in opposite directions there can be no single scalar that adjusts this genealogy to match one produced by a Wright-Fisher population. This is the term that is commonly used as *N_e_*, but may not always be the term that we actually want to consider in treating selective forces to understand protein evolution.

**Figure 1.**
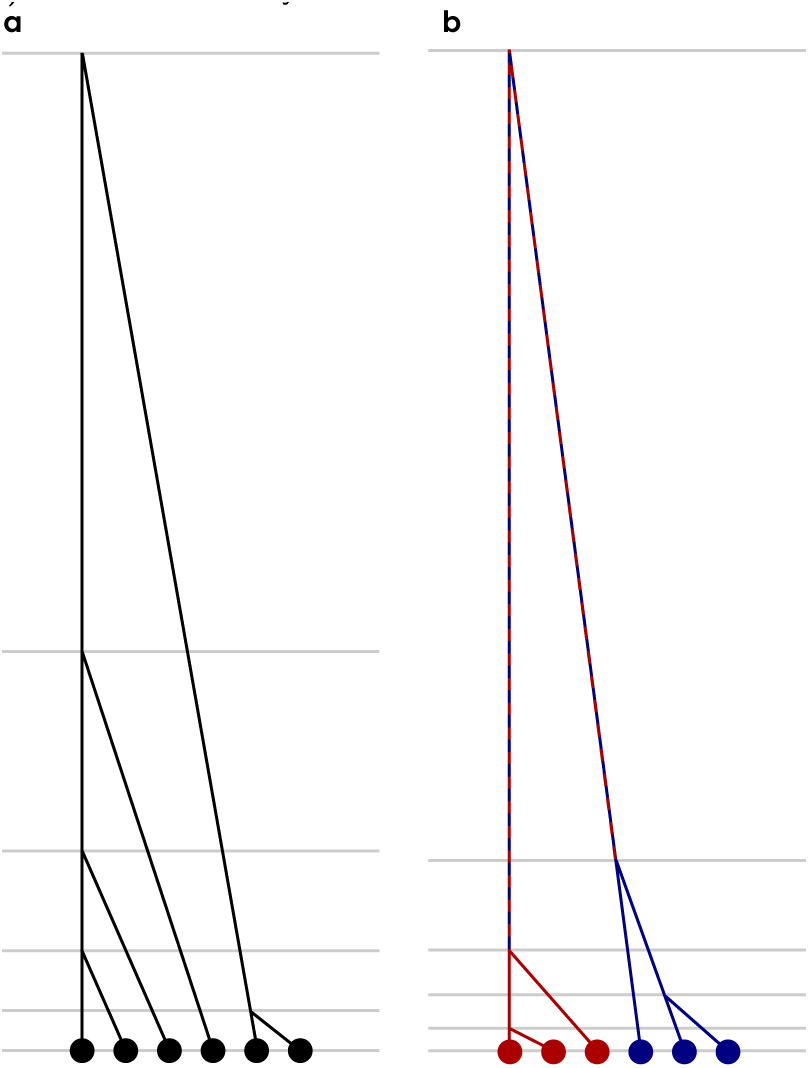
Example coalescent trees. Representative coalescent trees from (a) and unstructured population and (b) a population with two sub-populations with limited migration between them. Both trees have been scaled to the same total depth, but the distribution of expected coalescent times (grey lines) differ dramatically between the trees.

## Multiple Important Population Size Parameters

Removed from the cozy confines of the Wright-Fisher model, the simple multiplicative relationship between population size and the context-dependent conditional probability of acquiring a beneficial mutation, the genetic diversity, and the probability of fixation of a favored allele becomes more complex and requires different treatments of population size. If any individual born into a population has some chance *μ* of passing on a mutation, the population will acquire new mutations at the rate of *N_c_μ*. The population size scalar here does not depend on any properties of the relations among individuals within the population or how they came to be that way and is simply a reflection of the mutational target size. The loss of variation through random genetic drift, however, *does* depend on the nature of the population. A new neutral mutation enters a diploid population at frequency 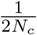. The expected time until fixation or loss of this mutation is 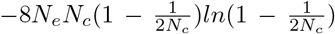, a function of both *N_e_* and *N_c_*. For mutations with an additive selection coefficient of *s*, the probability of fixation is (1 − *e*^−2*N_e_s*/*N_c_*^)/(1 − *e*^−4*N_e_*^*s*),or approximately 2*sN_e_*/*N_c_*, and is driven both by the intensity of selection on the particular variant and the ratio of the variance in reproductive output (as reflected in *N_e_*) and the mean reproductive output (as reflected in *N_c_*). In populations where *N_e_* cannot be properly defined, such as those with persistent subdivision, no general equation may suffice to capture all of these behaviors; mutations arise within each sub-population with probability proportional to the census size of the sub-population and then drift to loss or fixation according to a complex meta-population dynamic that may include considerable time spent fixed within some sub-populations and absent in others. While there will always be *some* probability distribution of time to fixation or loss of an arbitrary new mutant it is unlikely in these cases to be able to assign values *N_c_* and *N_e_* such that the distribution matches anything produced by a population that converges to the Kingman coalescent. These concepts need to be considered in the context of models for understanding selection in proteins.

## Practical Principles of Effective Population Size for Protein Evolution Studies

### Effective population size impacts the distribution of fitness effects

In addition to influencing the probability with which a mutation with a given selection coefficient fixes, it has been suggested that *N_e_* affects the distribution of selective effects itself [16, 3]. An important source of constraint on proteins arises from the requirement to fold stably into the correct structure [17, 8, 18]. The free energy of folding (Δ*G*) or stability of a given protein under stabilizing selection is determined by mutation-selection balance. This describes the equilibrium at which the rate of deleterious alleles being removed by selection equals the rate at which they arise due to mutation [19, 20]. As a result, proteins in nature are only marginally stable [21]. Because *N_e_* affects the efficacy of selection against deleterious, destabilizing variants, one might expect proteins to be more stable for organisms with large effective population sizes [4]. These differences in stability, then, are expected to impact the extent to which new mutations are stabilizing or destabilizing [22, 23, 24, 25]. In other words, the starting point affects the mutant’s fitness [26].

That the change in stability for new mutations, ΔΔ*G*, depends on Δ*G* is consistent with Fisher’s Geometric Model, where larger steps in phenotype space are more likely to land further from an optimum [27]. For example, at one extreme, if a protein were optimal, all possible mutations would be deleterious or neutral; at the other extreme, for a protein furthest from the optimum all possible mutations would be advantageous or neutral; at intermediate values, the relative fractions in the advantageous and deleterious categories are expected to vary. If all possible sequences are ranked by fitness as approximated by the fraction of protein folded into the active state, the probability density takes a form with highest mass in the middle and lowest at the extremes, stemming from results in statistical physics [28].

When a system is at equilibrium with constant Ne and constant selection, any changes that fix are largely neutral and independent of *N_e_*. A compensatory seascape where fitness fluctuates about an equilibrium driven by combinations of slightly deleterious changes and compensatory ones [29] is nevertheless possible for changes of small fitness effect. Compensatory evolution is expected to affect a larger fraction of changes at small Ne, where there is weaker selection against deleterious changes of equivalent magnitude and where mutations segregate for shorter periods of time before fixing and therefore have less chance of fixing together with interacting mutations [30, 31, 3]. The dynamics of fixation themselves are dependent upon *N_e_*. In small populations, mutations fix one at a time, creating a rugged fitness landscape. With larger populations, mutations and their compensatory changes may fix together due to stochastic tunneling [30]. This not only leads to less expected variance in Δ*G*, but also to faster neutral walks across sequence space, leading to stronger observed epistatic effects [32].

Further, Goldstein and Pollock [33] have implied that given the context dependence of fitness effects of mutations, termed epistasis, allowed and forbidden amino acid states play a greater role than the relative fitnesses of allowed states. This would result in a model where changes are either neutral or impermissible without a major contribution of positive selection to the compensatory process, corresponding to a substitution model with a neutral rate plus a shifting set of invariant substitutions. This is an interesting extension of a covarion model (involving transitions between variable and invariant states over time) [34] embedded in a substitution matrix. This relates to a model proposed by Usmanova and coworkers [35], where states transition between allowed and disallowed substitution states as a Markov process. The small differences in fitness values and corresponding selective coefficients for amino acid transitions may be an artifact of the substitution process model used in the simulations that generated these results, however. Future work will therefore be needed to evaluate if such a model actually explains protein sequence evolution well. The probability of fixation of an introduced mutation would be proportional to *N_e_* under such a model.

### Inferring fitness effects with mutation-selection models

Mutation-selection models allow the selective coefficients of amino acid changes to be inferred from sequence data. Given a set of observed codon substitutions over a phylogeny, we can assign each character state a fitness parameter based on the rate at which that state fixes once it has arisen. The relationship between the probability of fixation and inferred fitness of a character is based on the principles outlined above. The probability P of a substitution occurring therefore depends on *μ* (which in turn depends upon *N_c_*), the selective coefficient *s*, and *N_e_*. In contrast to many standard phylogenetic methods, the processes of mutation and selection can therefore be examined separately in this framework. This approach has recently become more commonly used in protein evolution studies, particularly with the availability more efficient computers [36, 37, 38, 39, 40]. It is of particular interest because it allows putatively adaptive or deleterious substitutions to be identified and separated from background noise in order to carefully characterize selection on proteins. Crucially, in order to obtain meaningful results from genomic data in this manner, we argue that a clear understanding of what is meant by *N_e_* is required and that *N_e_* and *N_c_* are treated distinctly.

By contrast, the most commonly used statistic to infer selection in coding sequences is the ratio of nonsynonymous to synonymous substitutions (dN/dS) [41, 42]. However, its limitations in terms of capturing evolutionary processes have become increasingly clear [33, 43]. For instance, at a given constant selection coefficient for amino acid replacements, the ratio will vary proportionally with the effective population size, and therefore cannot provide information about selection coefficients without additional inference of *N_e_*. Furthermore, this framework does not consider variable fitness effects conferred by different amino acid changes that can vary in terms of the severity of their impact on the protein structure. It also does not consider that exchanges between a given pair of amino acids may be favored in one direction, from a less fit to a more fit state, but disfavored in the other. These models are therefore consistent only with diversifying but not directional selection [33, 44]. On the other hand, the mutation-selection framework models the probability of introducing a particular kind of mutation by multiplying the per site, per individual mutation rate by the census population size. Once introduced, a mutation’s probability of fixation depends on the selective coefficient, the frequency at which it’s introduced into the population, and the population’s effective size. Applied to real populations with real sizes, structures, and complexities, the concept of the effective population size and its interpretation should be carefully considered. As populations deviate from abstract models, the notions of population size relevant for the introduction of a new mutation and its subsequent probability of fixation are not the same.

To describe the number of new mutations that may become available to a population as a forward looking measure, the relevant parameter is the current census population size. New mutations are introduced at frequency 1/2*N_c_* in a diploid population. The historical effective population size is important as a backward looking measure to describe the past effects of selection on mutations in a population [19]. In this framework, the probability of fixation of an amino acid replacement relative to a neutral variant is approximately described by Kimura’s diffusion equation [5], where the selection coefficient *s* and *N_e_* determine amino acid substitution properties. In this case, *N_e_* is the effective size of the population over a specific period of history and specific to lineages of phylogenetic trees where selection has been acting on the system in question. Where population sizes have changed rapidly, this may be a very different quantity than is relevant to describe the trajectories of existing or future variation. Furthermore, the relevant timescales for determining *N_e_* will not be constant across mutations. As selection acts more rapidly on variants conferring large fitness effects, the properties of these variants will reflect a more recent *N_e_* than neutral variants which will have experienced the effects of population parameters for a longer history. Generally, the probability of fixation of any particular mutation will depend on *N_c_* at the time it arose and the range of *N_e_* values during the period of time during which it existed at low frequency.

### Local Variation in *N_e_*

Demographic factors such as time-varying population sizes, inbreeding, or unequal sex ratios typically cause *N_e_* and *N_c_* to deviate from *N_c_* in a uniform manner across the entire genome. Other factors may create additional, local variation in specific parts of the genome. For instance, in species with heterogametic sexes, sex chromosomes have reduced effective population sizes *and* census population sizes relative to autosomes given their mode of transmission. Organellar genomes, such as those of mitochondria, show diverging locus-specific population sizes. While *N_c_* will be higher than that of the autosomal genome due to the high copy number of mitochondria in each cell, small inter-generational bottlenecks and transmission only through the female germ line leads to a *smaller N_e_* than seen in the autosomal genome. In accord with these predictions, rates of both heterozygosity and divergence have been shown to differ between autosomes and sex chromosomes (see, for instance, the fast-X effect [45]). Further, sex differences in the variance of reproductive success (as is a typical result of anisogamy) can lead to additional differences between sex chromosomes and autosomes in *N_e_* that are not reflected in differences in *N_c_*.

According to the population genetics perspective, loci linked to selected sites exhibit locally reduced effective population sizes through genetic hitchhiking, background selection, and Hill-Robertson interference [46, 47, 48, 49, 50]. For neutral loci, this is of little concern as the probability of fixation of a neutral mutation is 1/2*N_c_* and does not depend on *N_e_* at all. However, the effects of selection on linked sites drive up the variability in reproductive success at the focal site, increasing the rate of genetic drift and locally decreasing the depth of the local genealogies. This leads to regions of the genome experiencing higher rates of recombination exhibiting lower levels of linkage and therefore less local reduction in *N_e_* due to selection on linked sites ([51, 52, 53] and reviewed in [54]). As the fixation of a given mutation depends on the product of the selection coefficient and the ratio *N_e_*/*N_c_*, high rates of recombination can increase the relative rates of fixation of beneficial mutations (and decrease deleterious ones) compared to functionally equivalent mutations in regions of the genome with lower recombination rates or higher densities of constrained loci [55, 56, 57, 58].

Where it is feasible to model this local rescaling of the coalescent rate due to selection at linked loci with an explicit model based on the biological process it may be advantageous to our understanding of protein evolution to do so. To account for interference between linked sites, the fixation process can be modeled in linked blocks. Here, the probability of fixation for linked (but not functionally interacting) sites derives from the additive selective coefficients across the set of linked loci and a measure of *N_e_*, call it *N_e,demosraphic_*. This parameter scales the underlying Kingman coalescent tree of the population to account for all of the selective neutral processes that create deviations from the Wright-Fisher expectations. There are several possible solutions to implement the computation, including the use of Approximate Bayesian Computation based on simulation [59], inferring probable observed changes in a phylogenetic context, or approximating the effects of linkage through a sampling of the expected number of co-segregating changes coupled to a background distribution of *s* values. For longer evolutionary timescales, recombination can be incorporated into models of sequence evolution [60]. This would then cleanly separate *N_e_* and *s* in mutation-selection models and prevent parameter bleed [61] where *N_e_* and *s* would otherwise have co-linear effects on the shape of underlying local genealogies. At present, such approaches have only been implemented on a limited scale, and there exists ample scope to develop models that describe the processes driving protein evolution in a more elaborate manner. The identifiability of mutation-selection model parameters under different sets of assumptions is a current research topic [43].

## Further Issues Related to Model Realism

### Pitfalls in interpretation of *N_e_*

Great care must be taken in estimating population size parameters from one aspect of observed data with the goal of using it to infer or predict other evolutionary parameters. Though they may often be correlated, estimates of *N_e_* must be considered independently of measures of *N_c_*. If *N_e_* is estimated as a parameter from a model, it is not simply the mean *N_e_* over the branch, but exhibits more complex dynamics, and when derived from an observable statistic of a natural population that may not have a genealogy well-represented by a Kingman coalescent may not be serve as an appropriate estimate for *N_e_* in other contexts. Observations of segregating diversity can be used to estimate forward looking *N_e_* [62], but are best considered more strictly as estimates of future segregating diversity.

Unlike *N_e_* which is less a physical characteristic of a population than a descriptive one, Nc may in theory be unambiguously observed in nature. Outside of highly proscribed settings, however, actually doing so is difficult. Common methods include mark-and-recapture studies [63], plot sampling [64], or simply using body mass as a proxy for the inverse of the population size [65]. Proper estimation of *N_c_* is critical when mutation rates are estimated independently as a per base, per replication rate, as this rate is only useful in an evolutionary setting when scaled by the census population size.

Ratios of statistics involving common definitions of *N_e_* such as *dN*/*dS* can be particularly useful as backward looking estimators [66]. Where *dN* is approximately 2*μ_N_ sN_e_*/*N_c_* and *μ_S_*/(2*N_c_*), the *N_c_* terms cancel. With external estimates of the relative rates of non-synonymous and synonymous mutation (*μ_N_*/*μ_S_*) this gives us a direct estimate of 2*sN_e_*, a population-scaled selection coefficient often referred to as *S* in the protein evolution literature and *γ* by population geneticists. Any difficulties in correctly parameterizing or estimating *N_e_* in this case will result in complementary problems in estimating *s*, and *dN*/*dS* has been known to perform poorly as an estimator in certain cases [67, 66, 68, 69].

Genomic models not accounting for heterogeneity in *N_e_* will compound difficulties in disentangling *s* from *S* or *γ*. Differences in recombination rate between loci may be more challenging to account for than the reductions in *N_e_* that sex chromosomes experience. When *N_e_* is treated as a mechanistic parameter, it is important that it is accurately defined and reflects the effective population size without absorbing mis-specifications in the model (mis-parameterizing one parameter with effects that should be fit with a different parameter) [61]. This becomes particularly acute when *N_e_* is used in combination with *s*, a critical parameter for hypothesis testing in molecular evolution, for example when studying molecular adaptation.

A model that does not account for genomic heterogeneity in the mutation rate (associated with variables such as replication timing, heterozygosity and recombination rate) might also lead to incorrect inference of local *N_e_*. In order to describe protein evolution realistically, an appropriately complex model of selection that is robust to mutation, linkage, epistasis, and covarion-like behaviors that are induced is therefore likely necessary. Here, epistasis is conceptualized as a discrete biochemical process where a change at one position in a protein directly affects amino acid fitnesses at other positions in the same or other proteins, changing probabilities of fixing introduced mutations. When modeled as a site-independent process, this gives rise to covarion-like behavior, where rates of change at a position shift over time [70, 71]. In the simplest form ([34]), this involves a shift between a substitutable position and an invariant site.

## Conclusions

We have laid out how multiple distinct parameters associated with population size can jointly be used in mechanistic models of protein evolution. The goal of this discussion is to frame an understanding of population size that is cleanly separable from selection, and that has mechanistic meaning for the process of protein evolution. With this, models that capture the appropriate level of biological complexity to describe observed protein evolution data can be developed, enabling characterization of lineage-specific selective coefficients in comparative genomics.

## Author’s contributions

AP, CCW, and DAL all contributed to the concepts and text of this work.

## Acknowledgements

We thank Joanna Masel for discussions during the formulation of ideas that led to this work. This work was supported by NSF grant DBI-1515704.

